# How to build a mycelium: tradeoffs in fungal architectural traits

**DOI:** 10.1101/361253

**Authors:** Anika Lehmann, Weishuang Zheng, Katharina Soutschek, Matthias C. Rillig

**Author notes:** Corresponding author, Freie Universität Berlin, Institut für Biologie, Plant Ecology, Altensteinstr. 6, D-14195 Berlin, Germany.

## Abstract

The fungal mycelium represents the essence of the fungal lifestyle, and understanding how a mycelium is constructed is of fundamental importance in fungal biology and ecology. Previous studies have examined initial developmental patterns or focused on a few strains, often mutants of model species, and frequently grown under non-harmonized growth conditions; these factors currently collectively hamper systematic insights into rules of mycelium architecture. To address this, we here use a broader suite of fungi (31 species including members of the Ascomycota, Basidiomycota and Mucoromycotina), all isolated from the same soil, and test for ten architectural traits under standardized laboratory conditions.

We find great variability in traits among the saprobic fungal species, and detect several clear tradeoffs in mycelial architecture, for example between internodal length and hyphal diameter. Within the constraints so identified, we document otherwise great versatility in mycelium architecture in this set of fungi, and there was no evidence of trait ‘syndromes’ as might be expected.

Our results point to an important dimension of fungal properties with likely consequences for coexistence within local communities, as well as for functional complementarity (e.g. decomposition, soil aggregation).

## Introduction

The mycelium comprises the entirety of the hyphae of a fungus, representing its nutrient-capture and interaction interface, and the infrastructure for transport within the fungal individual. This structure is designed for a dynamic exploratory lifestyle with its ability to reconfigure, fragment and fuse, and represents the very essence of the fungal lifestyle^1^. Understanding how a mycelium is built is therefore of fundamental importance for gaining insight into fungal biology and ecology.

The initial development of the mycelium starting from germinating spores has been extensively studied (e.g.^2,3^), revealing some general hyphal growth patterns: emerging from a spore, a hypha extends at an exponential rate followed by a constant linear phase until the formation of a new branch is initiated; each new branch itself follows this exponential–linear phase pattern. Additionally, hyphae show negative autotropism and radial orientation away from the colony center^3^, eventually giving rise to the characteristic circular (in 2D) or spherical (in 3D) shape of ‘colonies’ or fungal individuals.

Mycologists have also examined the kinetics and branching behavior of fungi with a focus on a limited suite of traits and with a typical focus on mutants of the same species or a few species, mainly derived from fungal culture collections (e.g.^4-6^). These investigations revealed that fungal mycelia undergo changes in growth behavior due to differentiation. Studies on *Neurospora crassa* revealed that after approximately 22 h branching angles decrease while hyphal extension rate and diameters increase. Ultimately, the mycelium establishes a hierarchy in which hyphae of higher branching order have decreased hyphal growth rate and diameter in relation to the parental hyphae from which they emerged. As a consequence, the space-filling capacity of the mycelium increases^7^, leading to a maximum surface area while investing in a minimum of hyphal length^8^.

Apart from such studies of development, there is no systematic comparison of a range of architectural features, measured under the same, standardized laboratory conditions, on a larger set of fungi from a common ecological context. This is why we currently only have limited knowledge about tradeoffs governing mycelium architecture that could give insight on architectural ‘rules’. So far, insights into such tradeoffs come from studying single species or mutants, and this frequently has not resulted in a consensus. For example, the relationship between hyphal diameter and growth rate can be positive (e.g. *Bortrytis*) or neutral (e.g., *Mucor* strains)^9^. The same discrepancy holds true for the relationship between hyphal branching frequency (number of hyphal tips) and hyphal growth rate, which was examined in multiple studies focusing on *Neurospora* strains and mutants: some studies found a positive relationship^10^ and others did not^11^.

Recently, advantages of pursuing a trait-based approach in fungal ecology have been introduced^12,13^. One clear benefit of such an approach is to move beyond idiosyncratic comparisons of a few isolates to making comparisons using larger sets of fungal isolates, offering opportunities for more general inferences about mycelium architectural rules.

Here, we report on studies designed to collect mycelium architecture traits for a set of 31 saprobic fungal strains, containing members of the phyla Ascomycota, Basidiomycota and Mucoromycotina. Our goal was to uncover general ‘rules’ of mycelium construction by identifying tradeoffs among mycelium growth characteristics within this set of fungi, all isolated from the same soil.

## Materials and Methods

### Fungal strains

Fungal strains were originally cultured from Mallnow Lebus, a dry grassland in a nature conservation reserve (Brandenburg, Germany, 52° 27.778’ N, 14° 29.349’ E). A set of 31 fungal strains were isolated from soil samples as described in Andrade-Linares et al.^14^. Briefly, soils were diluted or washed to minimize spore abundance and increase the isolation of fungi derived from hyphae attached to soil particles^15,16^. For isolation a variety of media and antibiotics were used to target Ascomycota, Basidiomycota and Mucoromycotina while suppressing bacterial growth. Isolates were incubated at 22°C and were cultured on PDA. The fungal set comprised members of the Ascomycota (twenty strains), Basidiomycota (four strains) and Mucoromycotina (seven strains) (**Fig. 1a, Table S1**).

**Fig. 1.**
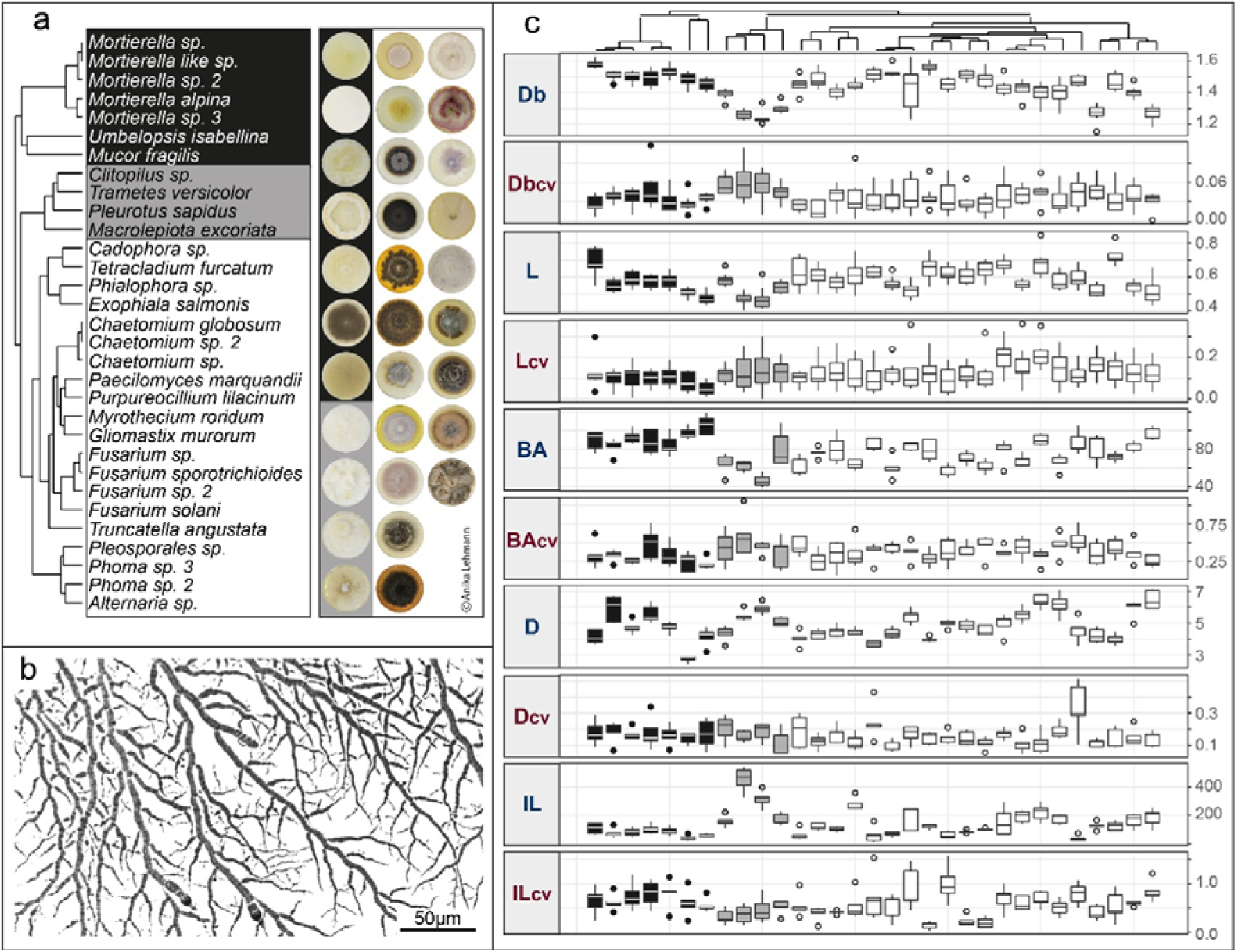
Overview of fungal strains, mycelial image and trait distribution. **(a)** Phylogenetic tree of the 31 fungal strains belonging to the phyla Ascomycota, Basidiomycota and Mucoromycotina with colony pictures. Pictures are from four-week old cultures grown on potato dextrose agar, and their order follows the order of the phylogenetic tree. Further information about phylogeny and accession numbers of the 31 strains are available in **Table S1**. **(b)** An example of mycelial pictures obtained from the setup introduced in the material and method section. Imaged strain is *Mucor fragilis.* **(c)** Tukey boxplot of the ten architectural trait variables, mean value and their coefficient of variation (CV) measured in this study: box counting dimension (unitless, Db with n= 8, Db_CV_ with n= 8), lacunarity (unitless, L with n= 8, L_CV_ with n=8), branching angle (in °, BA with n= 5, BA_CV_ with n= 5), hyphal diameter (in μm, D with n= 5, D_CV_ with n= 5), internodal length (in μm, IL with n= 5, IL_CV_ with n= 5). The boxplots represent 25th and 75th percentile, median and outlying points. Information about phylum affiliation is color-coded (black: Mucoromycotina, grey: Basidiomycota, white: Ascomycota).

### Architecture traits

We conducted two separate studies to collect architectural traits for the 31 fungal strains. All studies were performed *in vitro* with PDA as growth substrate. In the first study, we focused on measuring hyphal branching angle (BA), internodal length (IL) and diameter (D). For this, the fungal strains were grown on single concavity slides carrying 150 μl of PDA. We chose to reduce the concentration of PDA to 10% to obtain nutrient reduced growth medium for reduced mycelial density. This was necessary to be able to identify single hyphae in very densely growing fungi. To guarantee solidification of the medium, we added agar (Panreac AppliChem) to reach 15 g L^-1^ concentration. The growth medium was flattened by placing a cover slip on the liquid medium drop until it solidified. A pre-sterilized poppy seed carrying the target fungal strain was positioned in the center of the concavity. The slide was placed in a 9 cm Petri dish filled with a 5 mm layer of water agar to maintain high air humidity. Plates were sealed and stored at room temperature (22 °C) in the dark until the fungal colony covered half of the concavity area. For each fungal strain five slides were prepared and placed in separate Petri dishes. For the measurements, slides were examined under the microscope (Leica DM2500, bright field, 200x). Per slide, we randomly chose five hyphae as subsamples; for each of these hyphae we measured at the colony edge the last developed branching angle, the internodal lengths between this last and second-to-last branch and the hyphal diameters within this youngest internodal segment. For analyses, we used the image processing software ImageJ^17^. For each experimental unit, we calculated a mean value and a coefficient of variation (CV) from the subsample data. These represented two aspects of a trait: the average value and its variability. The trait data used in statistical analyses were the average of mean values and CVs of five replicates (i.e. n = 31).

In the second study, we investigated the complexity and the heterogeneity of fungal mycelia by applying fractal analysis – a technique used to assess self-similarity and space-filling capacity of fungal hyphae^18^. For this, we applied the same approach as in the first experiment but with eight replicates per fungal strain. At harvest, the slides were examined under the microscope (Leica DM2500, bright field, 200x) focusing on the outer 200 μm of the growing zone to investigate the “surface fractals”^19^. Camera (Leica DFC290) settings were chosen to generate grayscale photos with high contrast (background: white, hyphae: black; **Fig. 1b**). For each slide, we photographed three fields of view at the colony edge. These settings and further image processing in ImageJ^17^ and Adobe Illustrator (CS6, v.16.0.0) were necessary to guarantee comparable and unbiased photos that can be processed by image analysis software. First, photos were converted to 8-bit binary images in ImageJ and subsequently hyphae were skeletonized. For this, a thinning algorithm repeatedly reduced pixels from the edge of the target object until a one-pixel wide shape was reached^20^. In Illustrator, the skeletonized hyphae were reconnected and image artifacts excluded, if necessary. Line thickness was adjusted to mean diameter trait values derived from experiment one. The final processed images were loaded into the ImageJ plug-in “FracLac”^21^ to measure fractal dimensions. We chose box counting dimensions (Db) as a measure of structural complexity (i.e. the degree of detail or amount of parts a pattern consists of), and lacunarity (L) as a representative of structural heterogeneity (i.e. the gappiness or “rotational and translational invariance” in a pattern^21^). We used default settings but allowed for rotational orientations in analyses. Finally, subsample data were used to calculate CVs for box counting dimension and lacunarity. The subsample data were then merged to one mean and CV trait value per replicate. Additionally, we verified if implementing diameter data altered fractal dimension data by correlating skeletonized and adjusted diameter data for both box counting dimension and lacunarity (**Fig. S1**).

### Statistics

We analyzed the relationships between the ten architectural trait variables derived from 31 saprobic fungal strains represented by both mean value and coefficients of variation (CV) (n = 31). First, to evaluate fungal distribution in ten-dimensional trait space, we ran a principal component analysis using the function prcomp() in the package “stats” with z-transformed data. Significance of PC axes was determined via the function testdim()^22^ in the package “ade4”^23-25^. Only the first axis was significant, hence we included PC axis 1 and 2 in the visualizations without losing information from the excluded axes. Next, we conducted kernel density estimation to assess species occurrence probability following the procedure presented by Diaz et al.^26^. Briefly, we used kde() function of the “ks” package^27^ with unconstrained bandwidth selectors by implementing the function Hpi() on our first two PC axes. Using the function contourLevels() we estimated contour probabilities for 0.5 and 0.95 quantiles.

Second, to test for phylogenetic signal in our trait variables we used Moran’s I statistic, a measure for phylogenetic autocorrelation, as implemented in the package “phylosignal”. We accounted for phylogenetic relatedness among species (indicated by detected phylogenetic signals) by calculating phylogenetically independent contrast of our trait variables with the packages “picante”^28^ and “ape”^29^ using the functions pic() and match.phylo.data(). We evaluated if the assumptions of the Brownian motion model were satisfied by our data^30^. For that, we investigated the standardization of the contrasts via diagnostic regression tests to evaluate the relationship between absolute standardized contrasts and (i) the square root of their standard deviation^31^ and (ii) the node height (i.e. node age^32,33^). Identified influential nodes were excluded, following the threshold of absolute studentized residuals greater than 3^31,34^. To satisfy the Brownian motion model assumption, we used log transformed trait values und excluded two outlier nodes (node 49 and 61^35^).

Third, multiple pairwise correlations using Pearson’s rho were conducted and plotted with the function corrplot() in the eponymous package^36^. Analyses were done for original (non-transformed, n= 31) and phylogenetically corrected data (log-transformed, n= 28).

Fourth, we ran linear regressions and further investigated the relationships by quantile regression with the package “quantreg” (https://github.com/cran/quantreg). Under most ecological conditions, linear regressions tend to over- or underestimate relationships due to a focus on the mean of the response distribution. Especially in wedge-shaped data distributions, indicating unmeasured limiting factors, quantile regressions are more informative since they test the relationship between response and predictor variable at their maxima^37,38^. Both regression analyses were run on z-transformed data and model residuals were tested for homogeneity and normal distribution. Additionally, we ran multiple pairwise regressions for both original and phylogenetically corrected data to provide graphical information on data distributions of all trait combinations (see Fig. S2 and S3). These were generated by the function ggpairs() of the package GGally^39^.

All analyses were conducted in R (v. 3.4.1^40^) and plots were created with the graphics package ggplot2^41^ and its extension GGally.

## Results and discussion

### Trait expression

Overall, we found high variability among strains for all traits **(Fig.1**). The application of fractal dimensions on mycelium structure revealed that trait mean values of box counting dimensions (Db) ranged between 1.2 and 1.6, where a value of 1 represents a single unbranched hypha, and a value of 2 a complex space-filling mycelium. The most complex mycelium was found in the Mucoromycotina, while Basidiomycota had the most simply structured mycelia. For lacunarity (L), we found in our study that trait values ranged between 0.4 (Basidiomycota) and 0.7 (Ascomycota). With increasing trait value, the heterogeneity and hence gappiness of the mycelium increased. The investigation of hyphal features revealed that the branching angle (BA) varied substantially across fungal strains from 26 to 86° with Mucoromycotina having large angles and Basidiomycota rather small angles. For hyphal diameter (D) trait values ranged from 2.7 to 6.5 μm across the 31 strains where both extremes could be found in the Mucoromycotina. The length of the hyphal internodes (IL) showed considerable differences: Within Basidiomycota internodal lengths of 453 μm could be reached while in Mucoromycotina the lowest value of 40 μm was measured. Our values are within the range of previously reported architectural features of selected, individual saprobic filamentous fungi (e.g.^19,42-46^).

After establishing the trait database, we investigated the trait space generated by the collected fungal architectural features. To do this, we applied principal component analyses.

### PCA

For our 31 fungal strains, the sole significant first principal components accounted for 34% of the variability in the ten architecture traits (**Fig. 2a**). In this ten-dimensional trait space, the set of our 31 fungal strains occupied the whole PC plane with a clear separation of the three phyla across the plane. Considering the sole significant PC axis 1, Ascomycota assembled in the center flanked by Mucoromycotina on the left, driven by large branching angles and high mycelial complexity, and Basidiomycota on the right portion, primarily characterized by long internodes and wide hyphal diameters (**Fig. S4**). Across species, some clear correlations among traits became visible; hence, we further investigated the type and intensity of potential architectural tradeoffs for our fungal set.

**Fig. 2.**
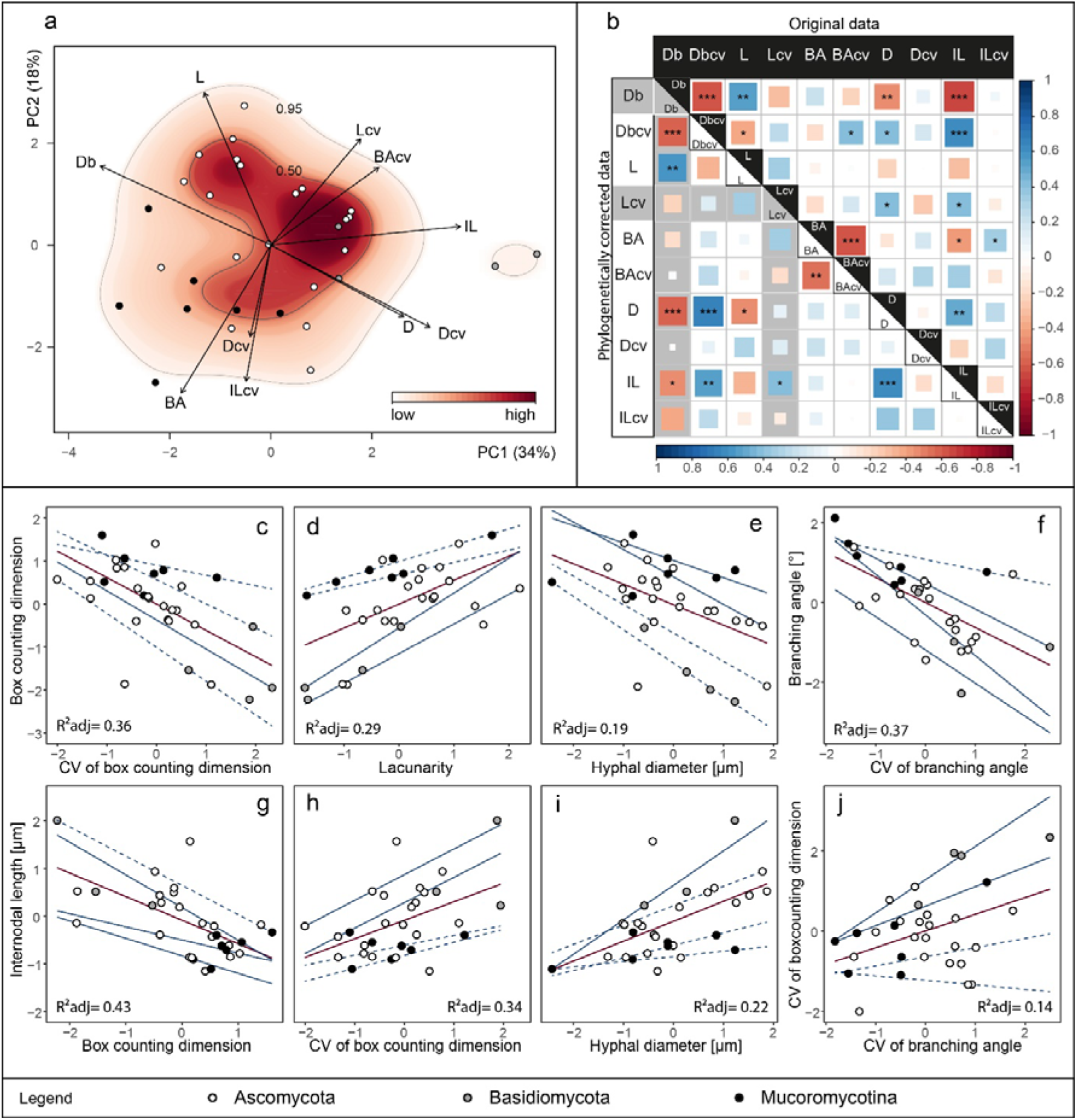
Outcomes of principal component analysis, trait correlation, linear and quantile regression of the investigated architectural traits. Analyses were conducted on trait data (n= 31). **(a)** Projection of the ordinated 31 fungal strains onto ten architectural trait variables (mean and CV): box counting dimension (Db), lacunarity (L), branching angle (BA), hyphal diameter (D), internodal length (IL) into two-dimensional trait space represented by principal component axis 1 and 2 (explaining 34 and 18% of variance, respectively). Arrows indicate direction and weight of trait vectors. Color gradient represents probability of species occurrence (white = low, red = high) in the trait space, with the contour lines denoting the 0.50 and 0.95 quantiles of kernel density estimation (see materials and methods section). **(b)** Correlation plot of five architectural trait variables and their coefficients of variation. The upper triangle displays original while the lower triangle represents phylogenetically corrected data correlations. Color gradient and square size are proportional to correlation coefficient (Pearson’s rho). Asterisks denote significance level: ^⋆⋆⋆^ < 0.001, ^⋆⋆^ <0.01, ^⋆^ < 0.05. In grey, we highlight trait combinations affected by detected phylogenetic signal (Table S2). **(c-j)** The eight strongest trait relationships for either original and/or phylogenetically corrected data. Red lines represent linear regression lines and blue lines quantile regression lines, while line type depicts significance of regression lines; solid lines p-value < 0.05, dashed lines > 0.05. Corresponding regression statistics can be found in Table S4. Adjusted R^2^ values correspond to linear regressions.

### Phylogenetic signal

For this, we first tested all ten traits for a phylogenetic signal to evaluate if the phylogenetic relatedness among fungal strains can influence any trait relationships we want to investigate. Applying Moran’s I statistics (**Table S2**), we found phylogenetic signals in Db (I= 0.16, p= 0.02) and L_CV_ (I= 0.13, p= 0.02). Hence, we needed to account for phylogenetic relations for these two traits among our 31 fungal strains by applying phylogenetically independent contrast in the following analyses.

### Tradeoffs

We found 14 trait pairs with significant correlations of which ten passed phylogenetic correction (**Fig. 2b, Fig.S2** and **S3**). The strongest correlations were detected between mycelium complexity and its coefficient of variation (Db - Db_CV_ in **Fig.2c**), mycelium heterogeneity, as measured by lacunarity, (Db - L in **Fig.2d**) and hyphal diameter (Db - D in **Fig.2e**), as well as between branching angle and its coefficient of variation (BA - BA_CV_ in **Fig.2f**). For internodal length, we detected relationships with mycelium complexity (IL - Db in **Fig.2g**), variability in mycelium complexity (IL - Db_CV_ in Fig.**2g**) and hyphal diameter (IL - D in **Fig. 2i**). Another strong correlation was found between the coefficients of variation of mycelium complexity and branching angle (Db_CV_ - BA_CV_ in **Fig.2j**). In addition, weak correlations were found for D and Db_CV_, D and L_CV_, as well as between IL and L_CV_, IL and BA (**Fig.S2** and **S3**). From these correlations we can deduce multiple rules for mycelium architecture.

For structural complexity (as represented by box counting dimensions) and branching angle we detected a negative relationship between their mean values and CVs (Db - Db_CV_ in **Fig. 2c** and BA - BA_CV_ in **Fig. 2f**). Thus strains exhibiting a high trait value for BA or Db are restricted to this high value, while strains with low values in these traits are capable of further adjusting these features.

Within strains, variability in mycelial complexity itself is determined by increasing internodal length (IL - Db_CV_ in **Fig. 2h**) and higher flexibility in branching angle measures (Db_CV_ - BA_CV_ in **Fig. 2j**). Thus, the degree of mycelial complexity can be modulated via branching patterns (e.g. distance between branches).

Considering the complexity – the space-filling capacity – of a mycelium, we found that more complex mycelia are more heterogeneously structured (Db - L in **Fig.2d**). Mycelia with high space-filling capacity tend to be rather heterogeneous in their structure, i.e. their mycelium is not uniformly complex but rather exhibits complex zones replaced by more simple mycelium structures towards the growing edge. At the colony edge, hyphae are confronted with new resources and environmental conditions for which a maximum of flexibility is likely advantageous. Furthermore, complex mycelia have smaller hyphal diameters (Db - D in **Fig. 2e**) and shorter internodal length (IL - Db in **Fig.2g**). A mycelium with long internodes is characterized by less branching and hence less space-filling. However, to be capable of growing long internodes the mycelium needs to improve its structural support, i.e. its tear-resistance. Long hyphae are at risk of fragmentation by shear-stresses^47^. To deal with this risk, hyphal cell walls can thicken and/or hyphal diameter can increase^5,48^. This is congruent with our finding that long internodes are linked with larger hyphal diameters (IL - D in **Fig.2i**).

### Conclusion

One of the most fundamental decisions a growing hypha has to make is when to branch. Thus, it is maybe not surprising that internodal length was a highly influential variable (aligned with PC axis 1, **Fig. 2**) in understanding the architecture of mycelia in trait space. This suggests that the trait internodal length is a main driver of mycelium architecture. Mycelia with short internodes can branch more frequently thus developing a more complex mycelium than those with long internodes. However, the capability of growing long unbranched hyphae enables the mycelium to more flexibly adjust their mycelial modules (see positive correlations between IL and Db_CV_, L_CV_) in response to environmental conditions.

It is interesting that there were no sharp boundaries in the sense of architectural ‘syndromes’ or clear groups of traits, but rather relatively gradual changes in trait values within the set of fungal isolates we examined. This illustrates the relative versatility of the mycelial growth form in evolutionary terms, at least in the peripheral growth zone of the fungus, which we examined here. We clearly show that there are limits to how a mycelium can be constructed, since some trait combinations are evidently not favorable (e.g. long internodes and small diameters). However, fungi have evidently otherwise filled the trait space within the constraints of such fundamental tradeoffs, even seen in a sample of 31 species. It will be interesting to compare our results to others sets of fungi once such data are available: our results suggest key parameters on which to focus.

Mycelial architecture is a fundamental property of filamentous fungi, governing the way these organisms explore their substrate. Using a set of fungi co-occurring in the same soil, we show that architectural features vary strongly and reproducibly among different isolates under the same laboratory conditions. It is therefore highly likely that such differences contribute to enabling coexistence within fungal communities^49^ by offering fungi different ways to forage and colonize the soil environment. On the other hand, such trait divergence can also mediate functional complementarity, for example in decomposition or soil aggregation^13^.

## Acknowledgements

This work was supported by the Deutsche Forschungsgemeinschaft (RI 1815/16-1). MCR additionally acknowledges an ERC Advanced Grant (694368).

